# *Plasmodium* secretion induces hepatocyte lysosome exocytosis and promotes parasite entry

**DOI:** 10.1101/618397

**Authors:** Kamalakannan Vijayan, Igor Cestari, Fred D. Mast, Elizabeth K.K. Glennon, Suzanne M McDermott, Heather S. Kain, Alyssa M. Brokaw, John D. Aitchison, Kenneth Stuart, Alexis Kaushansky

**Author notes:** Correspondence to: Alexis Kaushansky, 307 Westlake Ave, N. Suite 500, Seattle, WA 98109.

## Abstract

The invasion of a suitable host hepatocyte by *Plasmodium* sporozoites is an essential step in malaria infection. We demonstrate that in infected hepatocytes, lysosomes are redistributed away from the nucleus, and surface exposure of lysosomal-associated membrane protein (LAMP1) is increased. Lysosome exocytosis in infected cells occurs independently of sporozoite traversal. Instead, a sporozoite-secreted factor is sufficient for the process. Knockdown of the SNARE proteins involved in lysosome-plasma membrane fusion reduces lysosome exocytosis and *Plasmodium* infection. In contrast, promoting fusion between the lysosome and plasma membrane dramatically increases infection. Our work demonstrates new parallels between *Plasmodium* sporozoite entry of hepatocytes and infection by the excavate pathogen, *Trypanosoma cruzi* and raises the question of whether convergent evolution has shaped host cell invasion by divergent pathogens.

---

*Plasmodium* parasites, the causative agents of malaria, are transmitted to humans by the bite of infected female *Anopheles* mosquitoes. The sporozoite form of the parasite is deposited into human skin during a blood meal. Sporozoites are motile, and rapidly migrate through the skin to enter a capillary, which allows the parasite to travel to the liver. *Plasmodium* sporozoites have the capacity to transmigrate through cells using a process termed cell traversal (CT)(*1*). Once in the liver, sporozoites invade a single hepatocyte, form a parasitophorous vacuole (PV) and develop into a liver stage (LS) schizont, from which merozoites are released into the bloodstream and invade erythrocytes. The secretion of a multitude of sporozoite factors has been demonstrated to occur during motility, CT and, invasion, yet a precise role for most secreted factors remains undefined.

Membrane vesicle trafficking is fundamental to eukaryotic life and plays a regulatory role in nearly all cellular activities. Many intracellular pathogens target and subvert these trafficking events for their own benefit (*2, 3*). Previous work has demonstrated that the *Plasmodium* liver stage PV membrane co-localizes with late endosomes(*4*), lysosomes(*5*) and autophagic vesicles(*6*). When this association is initiated during *Plasmodium* life-cycle progression remains unknown. Moreover, the role of host vesicular trafficking processes in sporozoite entry of hepatocytes has not been explored.

We quantitatively surveyed the extent of co-localization between the parasite and five markers of endocytic compartments. Freshly isolated *Plasmodium yoelii* sporozoites were added to Hepa 1-6 cells. After 90 min, cells were fixed, stained, and visualized by 3D fluorescent deconvolution microscopy. We used antibodies against Early Endosome Antigen 1 (EEA1) and Ras-related protein 5 (Rab5) to mark early endosomes, Rab7a to mark late endosomes (LE), Rab11a to mark recycling endosomes and Lysosomal-associated membrane protein 1 (LAMP1) to mark LE/lysosomes. Sporozoites were labeled with an antibody against circumsporozoite protein (CSP) (Fig. 1a). Intensity based co-localization was used(*7*) to evaluate the extent of overlap between CSP and staining for each vesicular compartment (Fig. 1a, b). The Pearson’s correlation coefficient between CSP and LAMP1 was ~0.6, but staining did not significantly overlap between CSP and EEA1, Rab5, Rab7a or Rab11a (Fig. 1a, b). These data are consistent with earlier observations(*4, 5*).

**Fig. 1.**
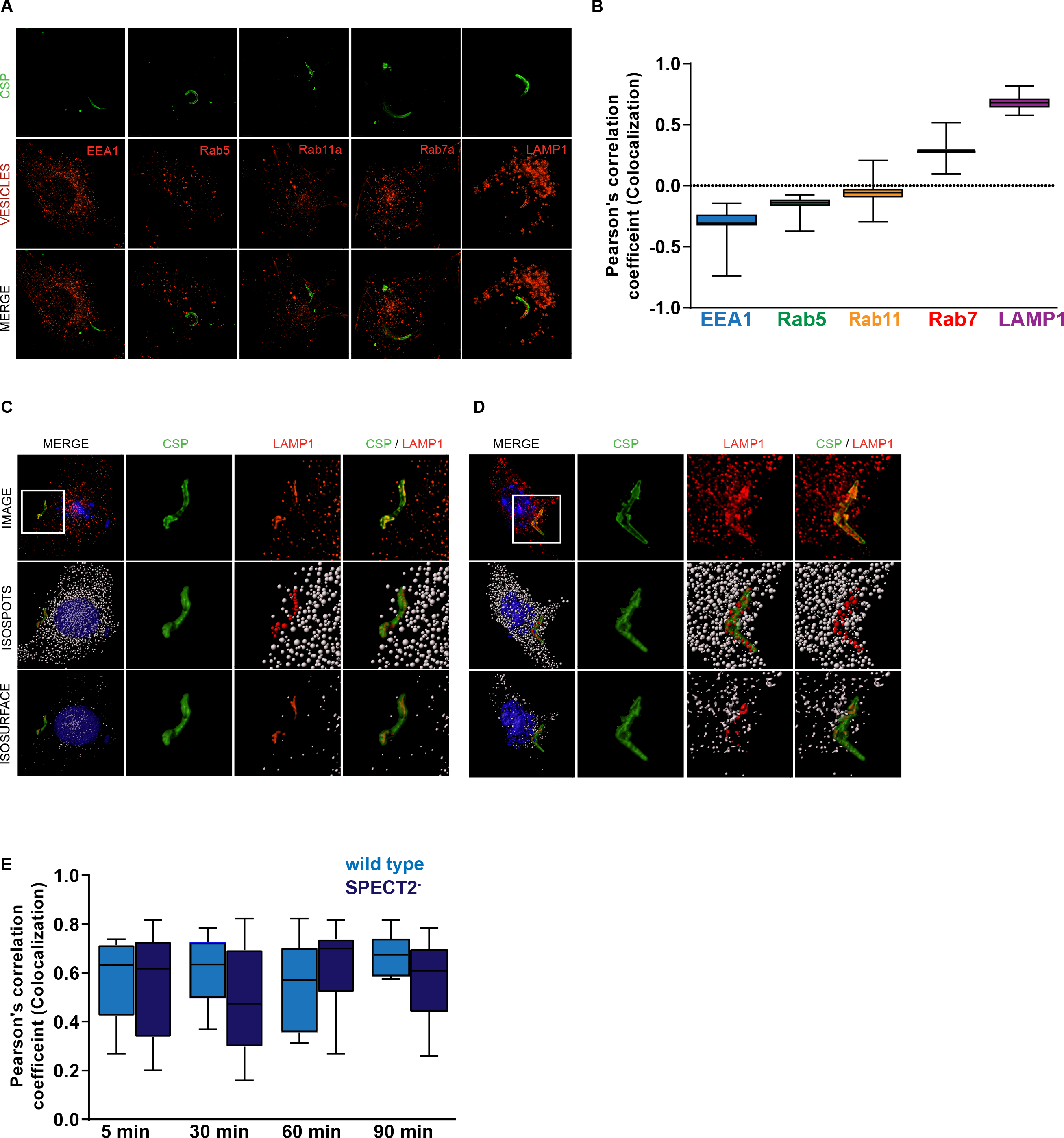
Interactions of *Plasmodium* sporozoites with early and late endocytic vesicles. **(a)** Hepa1-6 cells were infected with *P. yoelii* sporozoites for 90 min and processed for fluorescence microscopy using antibodies to EEA1, Rab5a, Rab11, Rab7a and LAMP1 (red) and *Py*CSP (green). Scale bar = 5 μm. **(b)** Pearson’s correlation coefficients were calculated for each endocytic vesicle channel with the sporozoite CSP channel of 25 different microscopic fields from 3 independent experiments. Hepa 1-6 cells were infected with *P. yoelii* sporozoites for 5 min **(c)** and 30 min **(d)** and processed for fluorescence microscopy using DAPI (blue) for DNA, Phalloidin (white) for actin visualization, antibodies to LAMP1 (red) for LE/lysosomes and CSP (green) for parasites. Isospot rendering for LE/lysosomes and isosurface rendering for LE/lysosomes, parasites, host cell nucleus and plasma membrane are shown. Red isospots represent LAMP1-positive structures co-localized with CSP. Magnified inset is 15 µm 𝑥𝑥 15 µm. **(e)** Hepa1-6 cells were infected with wild type or SPECT2^−^ *P. yoelii* sporozoites and fixed after 5, 30, 60, and 90 min. Intensity based colocalization was performed on at least 25 parasites per time point and Pearson’s correlation coefficients were calculated. Box and whiskers plot depict mean ± SD of three independent experiments.

We next assessed the kinetics of co-localization between sporozoites and LAMP1. Hepa1-6 cells were infected with *P. yoelii* sporozoites and fixed after 5 (Fig. 1c), 30 (Fig. 1d), 60 or 90 minutes (Fig. 1e). LAMP1 structures that co-localized with CSP were observed as early as 5 min and were elongated in the infected cells. These elongated LAMP1 structures were not observed in bystander or unexposed cells. Thus, the association between LAMP1-postive LE/lysosomes and sporozoites occurs during or very soon after infection and is maintained. We observed a very similar association between LAMP1 and CSP in the CT-deficient parasite, *Py*SPECT2^−^(*8*)(Fig. 1e, S1a and S1b). Our data are consistent with the hypothesis that lysosomes interact with the sporozoite during or very soon after entry, independently of CT.

Lysosomes are typically located in juxtanuclear regions of the cell under basal conditions but can be redistributed under times of stress or during infection(*9*). To evaluate lysosome localization during *Plasmodium* infection, we infected Hepa1-6 cells with *P. yoelli* sporozoites and fixed after 15 or 30 min (Fig. 2a). To assess the quantity and localization of lysosomes within infected and uninfected cells we defined LE/lysosomes as LAMP1-positive structures between 0.25 to 1 µm in diameter, corresponding to the typical size of LE/lysosomes within the mammalian cell. Hepa1-6 cells contained an average of ~450 LAMP1-positive structures, similar to measurements obtained by other groups(*10*). We defined perinuclear lysosomes as LAMP1-positive structures that were within a region surrounding the nucleus that was delineated by extrapolating the DAPI signal. In unexposed or mock treated samples containing material from the salivary glands of uninfected mosquitoes, ~85% of LE/lysosomes were perinuclear (Fig. 2a). In infected cells, lysosomes were slightly higher in number (Fig. 2b), but significantly less perinuclear (Fig. 2c). Interestingly, bystander cells, which were defined as being immediately proximate to the infected cell, also exhibited an increase in lysosome numbers and redistribution (Fig. 2b, c).

**Fig. 2.**
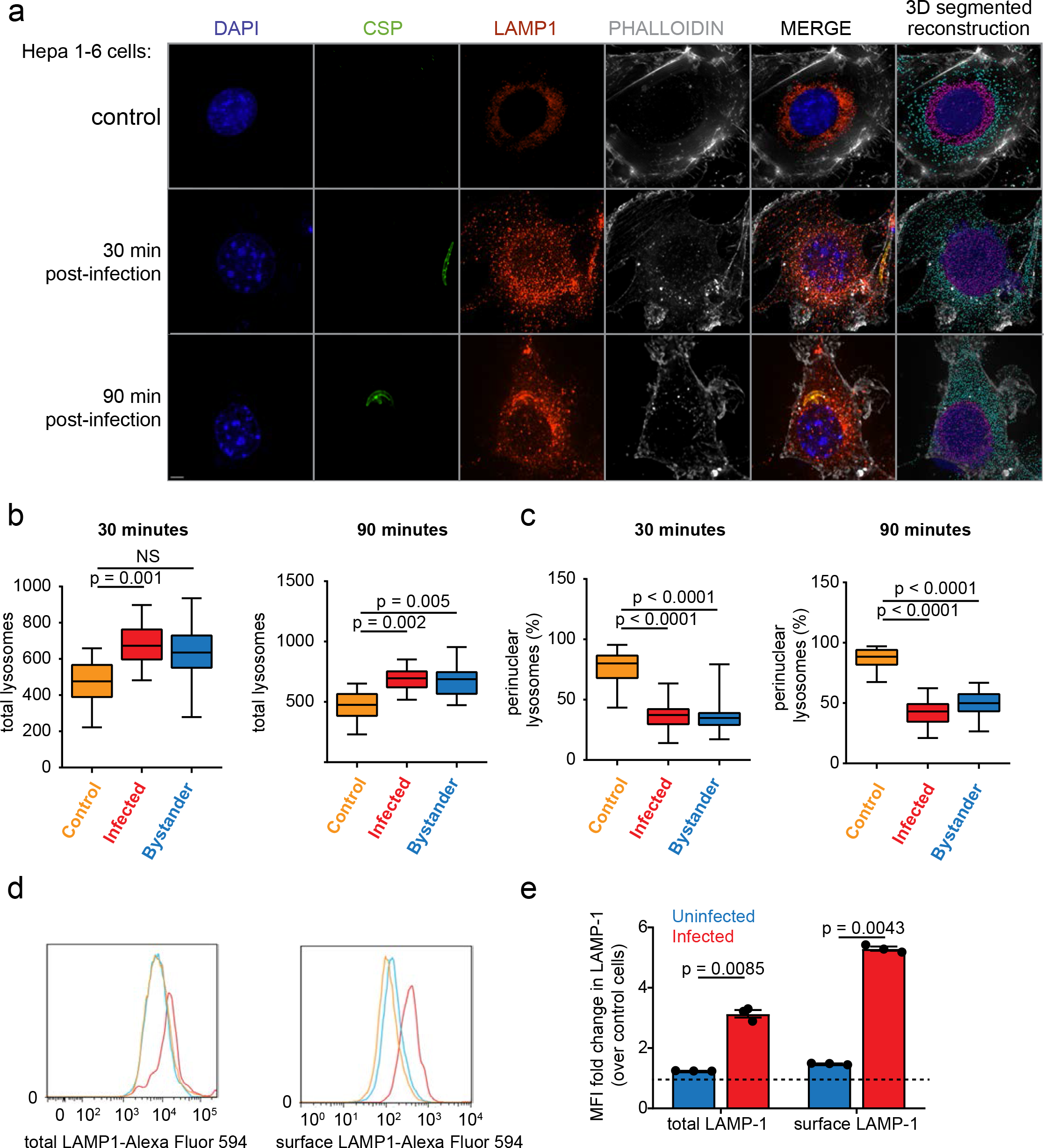
Sporozoite entry is associated with LE/lysosome redistribution during invasion. **(a)** Hepa1-6 cells were infected with *P. yoelii* sporozoites and fixed after 30 or 90 min. DNA was visualized with DAPI (blue), actin with phalloidin (white), LE/Lysosomes with anti-LAMP1 (red) and parasites with anti-CSP (green). Images are maximum intensity projections of the 3D dataset. Bar = 5 µm. The isospots corresponding to lysosomes away from the nucleus are cyan while perinuclear lysosomes are magenta. **(b and c)** Values represented in box and whiskers plots correspond to lysosomes from total and perinuclear area represented as mean ± SD of 25 different microscopic fields from 3 independent experiments. **(d)** Hepa1-6 cells were infected with *P. yoelii* sporozoites for 90 min and analyzed by flow cytometry using antibodies specific to LAMP1 and CSP. The histogram depicts the distribution of total and surface LAMP1 from infected, uninfected and unexposed control cells from one representative experiment. **(e)** Total and surface LAMP1 levels were compared between uninfected and infected cells as a fold change over control cells. The bar graph depicts the mean ± the SD of three independent experiments.

To assess the fate of redistributed lysosomes, we asked if there was evidence of LAMP1-vesicle fusion with the hepatocyte plasma membrane in infected cells. We infected Hepa1-6 cells with *P. yoelii* sporozoites and evaluated total and surface exposed LAMP1 (sLAMP1) by flow cytometry (Fig. 2d) and immunofluorescence microscopy (Fig. S2a). Both LAMP1 and sLAMP1 were elevated in infected cells compared to uninfected and unexposed cells (Fig. 2d). Interestingly, little if any impact on sLAMP1 was observed in uninfected cells, despite our earlier observation that lysosomes redistribute in these cells. These data suggest that lysosomes traffic away from the nucleus in infected and neighboring cells but undergo exocytosis only in infected cells. We observed a similar pattern of lysosome redistribution (Fig. S2b-d), and elevated levels of sLAMP1 (Fig. S2e, f) when we infected with *Py*SPECT2^−^. Together, these data suggest that lysosome trafficking and exocytosis are altered in infected hepatocytes, independently of CT.

The PV membrane (PVM) is critical for liver-stage development. Soon after productive infection, parasite factors, including upregulated in infectious sporozoites 4 (UIS4), are translated and trafficked to the PVM (*11*). We infected Hepa1-6 cells with wild type *P. yoelli* sporozoites and fixed samples 3 h after infection. LS parasites with an intact PVM were distinguished by positive CSP and UIS4 staining, and LS parasites positive for CSP but negative for UIS4 were defined as having a non-existent or compromised PVM. We observed co-localization of LAMP1 with parasite markers in both cases (Fig. S2g) suggesting that lysosomal contents are associated with all intracellular parasites, independently of the status of their PVM.

Lysosomes have been previously demonstrated to play a role in *Trypanosoma cruzi* host cell entry(*12*). Specifically, a portion of *T. cruzi* parasites utilize a lysosome-mediated event to enter the host cell(*12, 13*). In contrast, *Toxoplasma gondii*, an apicomplexan parasite closely related to *Plasmodium*, sequesters host lysosomes to the vacuolar space (*14*), but is not thought to use lysosomes to facilitate host cell entry. To elucidate how the invasion of *Plasmodium* parasites compares to these two disparate systems, we infected Hepa1-6 cells with *P. yoelii* sporozoites, *T. gondii* tachyzoites or *T. cruzi* trypomastigotes and assessed infection and sLAMP1 by flow cytometry after 90 min. Cells infected with *P. yoelii* or *T. cruzi*, but not *T. gondii*, exhibited increased sLAMP1 (Fig. 3a, b).

**Fig. 3.**
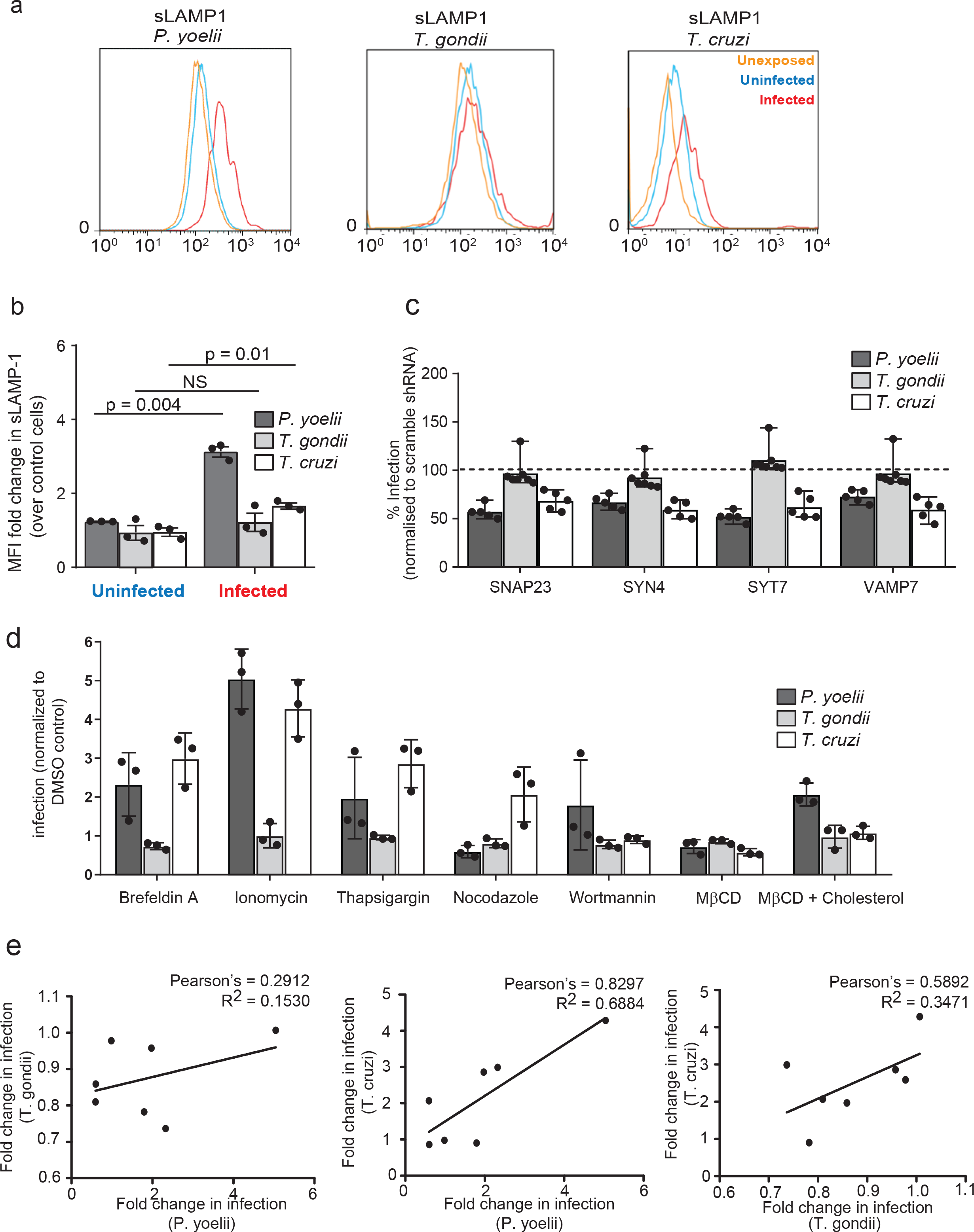
The role of lysosome exocytosis varies across species of intracellular parasites. **(a, b)** Hepa1-6 cells were infected with *P. yoelii* sporozoites, *T. gondii* tachyozoites or *T. cruzi* trypomastigotes for 90 min and analyzed by flow cytometry. The histogram shows surface LAMP1 from infected, uninfected and unexposed control cells. Surface LAMP1 is expressed as fold change over uninfected cells. The bar graph displays the mean ± SD of three independent experiments. **(c)** Hepa1-6 cells were transduced with shRNA lentiviruses against SNAP23, SYN4, SYT7, VAMP7 or a scrambled control and challenged with *P. yoelii* sporozoites, *T. gondii* tachyozoites or *T. cruzi* trypomastigotes for 90 min. The bar graph displays the infection rate after knockdown of each transcript of interest normalized to scramble shRNA cells, indicated by dashed line (n=5 for *Plasmodium* and *Trypanosoma*; n=7 for *Toxoplasma* infection; mean ± SD). **(d)** Hepa1-6 cells were incubated with or without the indicated compound for 15 min, washed and then infected with the indicated parasite for 90 min. The bar graph represents mean ± SD of three independent experiments. **(e)** Pearson’s correlation coefficients were calculated from the data in **(d)** for each pairwise combination of infections.

Fusion between lysosomes and the plasma membrane is mediated by the SNARE complex, which includes Synaptotagmin VII (SYT7), Syntaxin 4 (SYN4), vesicle associate membrane protein 7 (VAMP7) and Synaptosomal-associated protein 23 (SNAP23)(*15*). We knocked down each factor in Hepa1-6 cells using lentivirus-encoded shRNAs and observed decreased levels of transcript (Fig. S3A) and reduced sLAMP1 (Fig. S3b)(*15*). We infected each knockdown line with *T. gondii*, *P. yoelii* or *T. cruzi* parasites. Knockdown of each member of the SNARE complex significantly reduced *P. yoelii* and *T. cruzi* infections, but not *T. gondii* infection (Fig. 3c). Thus, *Plasmodium* sporozoites may use similar host cell machinery to infect as the excavate parasite *Trypanosoma cruzi*. In contrast, the apicomplexan parasites, *Plasmodium* and *Toxoplasma*, do not share this feature.

Genetic knockdowns can sometimes lead to compensatory changes that produce off-target effects. To partially circumvent this, we evaluated the impact of a range of small molecules that modulate lysosome exocytosis (Table S1). We treated Hepa1-6 cells with each compound for 15 min, washed the cells, and then infected with *P. yoelii* sporozoites, *T. gondii* tachyzoites or *T. cruzi* trypomastigotes for 90 min. Molecules that increase lysosome exocytosis or redistribution (Ionomycin, Thapsigargin, Brefeldin A, Table S1 and Fig. S3C), significantly increased *P. yoelii* and *T. cruzi* infection (Fig. 3d). In contrast, molecules that reduced lysosome exocytosis or redistribution (Nocodazole, MβCD) diminished *P. yoelii* and *T. cruzi* infections. No inhibitors substantially altered *T. gondii* infection (Fig. 3d). Overall, changes in *P. yoelii* and *T. cruzi* infections were tightly correlated (Pearson Correlation Coefficient = 0.8297) (Fig. 3e), while other pairwise comparisons were less correlated. Our data suggest that *T. cruzi* and *P. yoelii*, but not *T. gondii*, rely on a lysosome-mediated mechanism to enter the host cell. The extent of these parallels, and ways in which the entry strategies diverge, remains an important area for further investigation.

Lysosome exocytosis is induced by the soluble *T. cruzi* factor *Tc*gp82(*10*). We treated sporozoites with FBS to induce secretion and then collected supernatants. We exposed Hepa1-6 cells to this sporozoite-derived, secretion-enriched supernatant at different sporozoite:hepatocyte ratios for 90 min. Cells were monitored for lysosome redistribution by 3D fluorescence microscopy (Fig. 4a, b, S4a) and sLAMP1 by flow cytometry (Fig. 4c). Treating cells with even low quantities of secretion-enriched supernatant, but not heat-inactivated supernatant, promoted lysosome redistribution (Fig. 4b), but sLAMP1 was induced in a dose-dependent manner (Fig. 4c). Therefore, sporozoite-induced lysosome redistribution is impacted by different factors, or the same factors at different levels than lysosome exocytosis. These results are consistent with a model where two separate secretion-mediated events induce hepatocyte lysosome redistribution and lysosome exocytosis (Fig. 4d). Taken together, our data suggest a role for lysosome exocytosis in hepatocyte entry of sporozoites, independently of CT or the presence of the PVM.

**Fig. 4.**
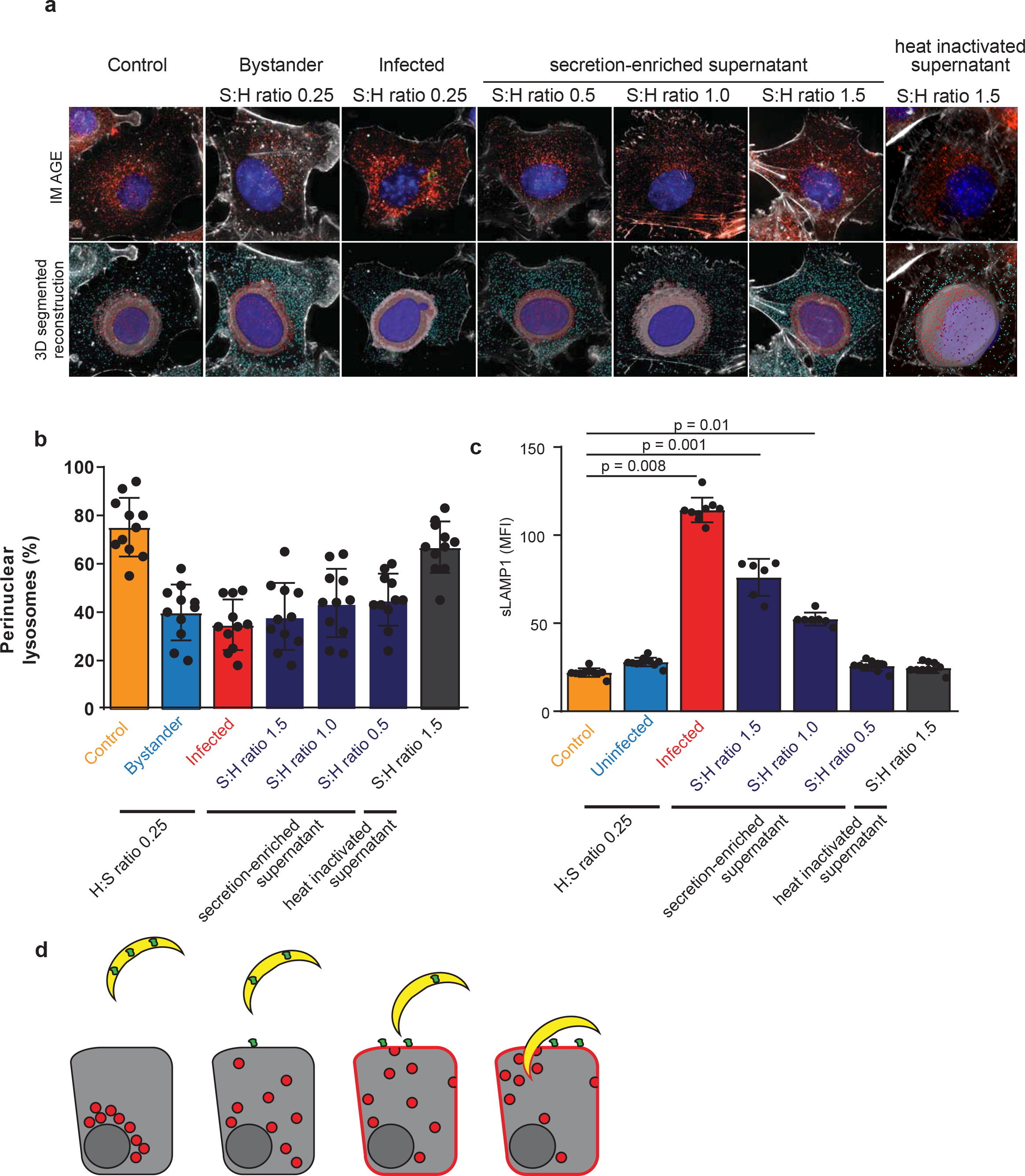
Sporozoite secretory factor(s) contribute to lysosome redistribution during invasion. **(a)** Hepa1-6 cells were infected with *P. yoelii* sporozoites or treated with sporozoite secretion-enriched supernatants at different sporozoite:hepatocyte (S:H) ratios. After 90 min, cells were processed using DAPI (blue) for DNA, phalloidin (white) for actin visualization, antibodies to LAMP1 (red) for LE/lysosomes and CSP (green) for parasites and displayed as maximum intensity projections. Bar = 5 µm. The isospots corresponding to lysosomes away from the nucleus and perinuclear lysosomes were depicted in cyan and magenta, respectively. **(b)** Values represented in bar graphs correspond to percentage of perinuclear lysosomes identified in (a). Data represents the mean ± SD of at least 10 different microscopic fields per condition from three independent experiments. **(c)** Hepa1-6 cells were infected with *P. yoelii* sporozoites or exposed to sporozoite secretion-enriched supernatants. Surface LAMP1 was analyzed by flow cytometry. Values represent the mean ± the SD of three independent experiments. **(d)** Model of *Plasmodium* sporozoites promoting lysosome exocytosis. Sporozoites are depicted in yellow, hepatocytes in grey, lysosomes in red, and a *Plasmodium*-derived secreted factor in green.

A growing collection of evidence suggests that parasites that differ only slightly in genetic makeup can exhibit drastically altered host cell tropism. For example, *Plasmodium* species rely differentially on host proteins CD81 and SRB1 for entry(*16–18*) and this relationship cannot be predicted by evolutionary similarity alone(*19*). Here, we demonstrate that lysosome-related alterations impact *P. yoelii* and *T. cruzi* infections similarly, but have no effect on the apicomplexan parasite, *T. gondii*. These observations raise the question of how quickly pathogens can evolve host cell tropism, and whether the similarities we observe are sculpted by convergent evolution. Systematic investigation of mechanisms of host cell invasion, across many pathogens, with well-defined evolutionary relationships, will allow us to obtain a better understanding of the major influences that shape host cell engagement over evolutionary time.

## Acknowledgements

We thank Marilyn Parsons for the RHΔHXGPRT *T. gondii* strain. We thank the Center for Infectious Disease Research/ Seattle Children’s Research Institute vivarium staff for their work with mice. All work was done according to IACUC procedures and protocols. This work was supported by National Institutes of Health grants R01GM101183 (AK), 1K99/R00AI111785 (AK), R01AI014102 (KS, IC and SM), P41 GM109824. (JDA), T32 Post-Doctoral Fellowship AI07509 (EKKG), a W.M. Keck Foundataion award (AK), Science and Engineering Research Board and INDO-US Science and Technology Forum – Overseas Post-Doctoral Fellowship (KV). FDM is a postdoctoral fellow with the Canadian Institutes of Health Research.

## Author Contributions

KV, EKKG, IC, SM, HSK and AMB performed experiments. KV, IC and FDM analyzed data. JDA, KS and AK supervised the research. KV and AK wrote the paper with input from all other authors.

## Conflict of Interest

The authors declare that they have no conflict of interest.

## Data availability

The data supporting the findings of this study are available within the paper and its Supplementary Information.

