## Supplementary files for "*Plasmodium* secretion induces hepatocyte lysosome exocytosis and promotes parasite entry"

#### Materials and Methods

##### Cell lines and culture

Hepa1-6 cells were obtained from American Type Culture Collection. Cells were maintained in DMEM-Complete Medium (Dulbecco's modified eagle medium (Cellgro, Manassas, VA), supplemented with 10% FBS (Sigma-Aldrich, St. Louis, MO), 10000 IU/ml penicillin/100 mg/ml) streptomycin (Cellgro), 2.5 mg/ml fungizone (HyClone/Thermo Fisher, Waltham, MA) and 4 mM L-Glutamine (Cellgro). Cells were split 2-3 times weekly. All experiments were performed using Hepa1-6 cells that were passaged between 4 and 20 times after purchase from ATCC.

##### Mosquito rearing and sporozoite production

For *P. yoelii* sporozoite production, female 6–8-week-old Swiss Webster mice (Harlan, Indianapolis, IN) were injected with blood stage *P. yoelii* (17XNL) parasites to begin the growth cycle. Animal handling was conducted according to the Institutional Animal Care and Use Committee-approved protocols. *Anopheles stephensi* mosquitoes were allowed to feed on infected mice after gametocyte exflagellation was observed. Salivary gland sporozoites were isolated using a standard protocol at day 14 or 15 post-blood meal. The sporozoites were activated with 20% FBS and spun at 1000g to remove debris from salivary gland. Spin sporozoites at 15,000 × g for 4 min at 4°C to pellet and resuspend in the desired volume of complete medium.

##### *T. gondii* production

*T. gondii* strain RHΔHXGPRT Gra2:GFP, Tub:βgal, a kind gift from Marilyn Parsons (CIDR, Seattle), was maintained by continual cycling through human foreskin fibroblasts (HFF). For infections, parasites were lysed from HFFs by passing 2× through a 27-gauge needle and then counted on a hemocytometer.

##### ***T. cruzi* production and labeling**

Tissue culture-derived trypomastigotes from the *T. cruzi* Cl Brener strain were obtained by weekly passage in confluent monolayers of Hepa1-6 cells at 37 ˚C and 5% $CO_2$, in DMEM medium supplemented with 10% FBS. Motile trypomastigotes were obtained from the supernatant and purified as previously described (*1*) and stained with 5(6)-Carboxyfluorescein diacetate N-succinimidyl ester (Sigma Aldrich) as suggested by manufacturer's protocol.

##### **shRNA-mediated gene knockdown**

MISSION shRNA vectors for SNAP23, VAMP7, SYT7 and SYN4 were obtained from Sigma Aldrich (St. Louis, MO). Non-replicating lentiviral stocks were generated by transfection of HEK293-FT cells. $4 \times 10^6$ HEK293-FT cells were plated on poly-L-lysine coated dishes to achieve 70-80% confluency at time of transfection. Approximately 24 h after plating, transfection mixtures were prepared by mixing 20 µL Polyethylenimine MAX (Polysciences Inc, Warrington, PA) prepared at 1 mg/ml, together with 4.75 µg of shRNA construct or a scramble shRNA control, 1.5 µg viral envelope plasmid (pCMV-VSV-G), and 3.75 µg viral packaging plasmid (psPax2). After incubating for 10 min at room temp in DMEM, transfection complexes were added drop-wise to cells. After overnight incubation, cells were washed to remove transfection mixtures and were fed with 10 mL fresh media. Lentivirus-containing supernatant was harvested 36 hours later, passed through 0.45 µm syringe filters, and either used immediately for transduction or stored at -80 ˚C.

To induce knockdown of candidate SNARE proteins, Hepa1-6 cells were transduced with lentiviral supernatants in 6-well plates at a cell density of $1 \times 10^6$ per well. At time of plating, cells were transduced with 1 mL of supernatant in the presence of 0.5 µg/mL polybrene (Sigma Aldrich St. Louis, MO). In order to select for cells with stable integration of shRNA transgenes, supernatant was replaced with complete media with the addition of 2 µg/mL puromycin 24 h post-transduction, and cells were selected for at least 5 days prior to experiments.

**Validation and quantification of shRNA mediated knockdown**

Total RNA was extracted using TRIzol reagent according to the manufacturer's procedure (Invitrogen). cDNA synthesis was performed using the Thermo Scientific RevertAid RT Kit according to the manufacturer's instructions (Thermo Scientific). For quantitative PCR (qPCR) a standard curve was generated using 1:4 dilutions of a reference cDNA sample for PCR amplification of all target PCR products using target specific primers (Table S1). The values of each transcript were normalized to mouse GAPDH. Experimental samples were compared to this standard curve to give a relative abundance of transcript.

**Infection assays**

$5 \times 10^5$ Hepa1-6 wild type cells or knockdowns cells were seeded in each well of a 24-well plate (Corning) and infected with *P. yoelii* sporozoites at a multiplicity of infection (MOI) = 0.25, *Toxoplasma gondii* tachyzoites (MOI = 0.5) or *Trypanosoma cruzi* trypomastigotes (MOI = 5) for 90 min. For small molecule treatment experiments, Hepa1-6 cells were treated with or without 10 µM ionomycin, 400 nM thapsigargin, 10 µM brefeldin A, 10 µM nocodazole and 5mM methyl-β-cyclodextrin (MβCD) for 15 min, washed and infected for 90 min. For cholesterol replenishment, MβCD treated cells were washed and incubated with 1 mM cholesterol for 15 min before parasite addition. After 90 min of infection, cells were harvested with accutase (Life technologies) and fixed with Cytoperm/Cytofix (BD Biosciences). Cells were blocked with Perm/Wash (BD

Biosciences) + 2% BSA for one hour at room temperature then stained overnight at 4 ˚C with primary antibody. Cells were washed three times with PBS then stained for one hour at room temperature with secondary antibodies. The cells were then washed and resuspended in PBS + 5 mM EDTA. Infection rate was measured by flow cytometry on an LSRII (Becton-Dickinson) and analyzed by FlowJo (Tree Star).

For the evaluation of surface expression of LAMP-1, cells were detached using Accutase (Sigma) and were incubated with a polyclonal antibody to LAMP1 (DSHB) in medium + 2% BSA for 30 minutes in ice, fixed with 3.7% paraformaldehyde, permeabilized with 0.01% Triton X-100. Cells were then stained with monoclonal antibody to *P. yoelii* Circumsporozoite protein (CSP) conjugated to AlexaFluor 488 (Life Technologies) at 1:500, *T. gondii* P30 mouse monoclonal antibody at 1:1000 (Novus Biologicals). Cells were washed three times with PBS then stained for one hour at room temperature with secondary antibodies. The cells were washed and suspended in PBS+5 mM EDTA. Infection rate and surface expression of LAMP1 was measured by flow cytometry on an LSRII (Becton-Dickinson) and analyzed by FlowJo (Tree Star).

**3D Fluorescence Microscopy**

For imaging experiments, Hepa1-6 cells were plated in 8 well chamber slides (Labtek) and infected with *P. yoelii* wild type or SPECT2$^{-}$ sporozoites. Cells were fixed with 10% formalin (Sigma) at defined timepoints after infection (5, 30, 60, or 90 min), permeabilized with Trition X-100, and stained with rabbit anti-EEA1 (CST), rabbit anti-Rab5 (CST), rabbit anti-Rab7 (CST), rabbit anti-Rab11 (CST), rat anti-LAMP1 (DSHB), goat anti-UIS4 (SCIGEN) and mouse anti-CSP antibodies. Nuclei were stained with DAPI (Vectashield) and Alexafluor 647-phalloidin (Life technologies) was used for actin visualization. Images were acquired with a 100× 1.4 NA objective

(Olympus) on a DeltaVision Elite High Resolution Microscope (GE Healthcare Life Sciences). The sides of each pixel represent 64.5 × 64.5 nm and z-stacks were acquired at 300 nm intervals. Approximately 20-30 slices were acquired per image stack. For deconvolution, the 3D data sets were processed to remove noise and reassign blur by an iterative Classic Maximum Likelihood Estimation widefield algorithm provided by Huygens Professional Software (Scientific Volume Imaging BV, The Netherlands).

**Image analysis and quantification**

Imaris software (Bitplane) was used to obtain 3D reconstructions of the fluorescence microscopy image stacks and quantification of lysosomes in x-y-z coordinates. Deconvolved images of immunostained cells stained with anti-LAMP1 (lysosomes) anti-CSP (sporozoites) and phalloidin (actin) with DAPI (nucleus) were processed, thresholded and segmented by Imaris software, to render isospots and isosurfaces from the fluorescence signal (*2*). Pahlloidin channel was used for the 3D reconstruction of the cells and only the sporozoites encased inside the 3D phalloidin structure are considered as invaded sporozoites and proceeded further. Isosurfaces were constructed by extrapolating the DAPI signal to the local minima in order to define a perinuclear region where lysosomes can be differentially counted. Using a mask tool, all LAMP1 signal outside this nuclear/perinuclear isosurface was suppressed, allowing addition of a new LAMP1 fluorescence channel corresponding exclusively to LAMP1 localized in perinuclear area. After image processing, we obtained an unmasked LAMP1 signal corresponding to total lysosomes and a masked LAMP1 signal localized to the perinuclear region, corresponding to perinuclear lysosomes. Isospots were constructed based on these two classes of LAMP1 signal, which allowed the quantification of total and perinuclear lysosomes per cell for each image-stack. The surface segmentation function of Imaris was used to identify the cell boundary using phalloidin-signal.

Intensity based co-localization was performed by creating region of interest (ROI) specific to the sporozoite structure in the CSP channel using the Imaris isosurface module. Pearson's correlation coefficient for co-localization analysis of endocytic vesicles and CSP was performed in the ROI using the Imaris co-localization module.

**Generation of sporozoite supernatant**

Salivary gland sporozoites were isolated using standard protocols. Sporozoites incubated with 20% FBS for 20 min at RT and spun at 13,0000 X *g* for 4 min at 4 ˚C. The supernatant is collected and again spun at 13, 0000 X g for 4 min at 4 ˚C to ensure the preparation is free of intact sporozoites. Hepa1-6 cells were exposed to supernatants at different sporozoite to hepatocyte ratio for 90 min. Cells were washed and subjected to 3D immunofluorescence microscopy or flow cytometry as described previously.

**Statistical analysis**

p-values were determined in GraphPad Prism 8 software using two tailed end t-test for samples with unequal variance.

### Supplementary Figure 1

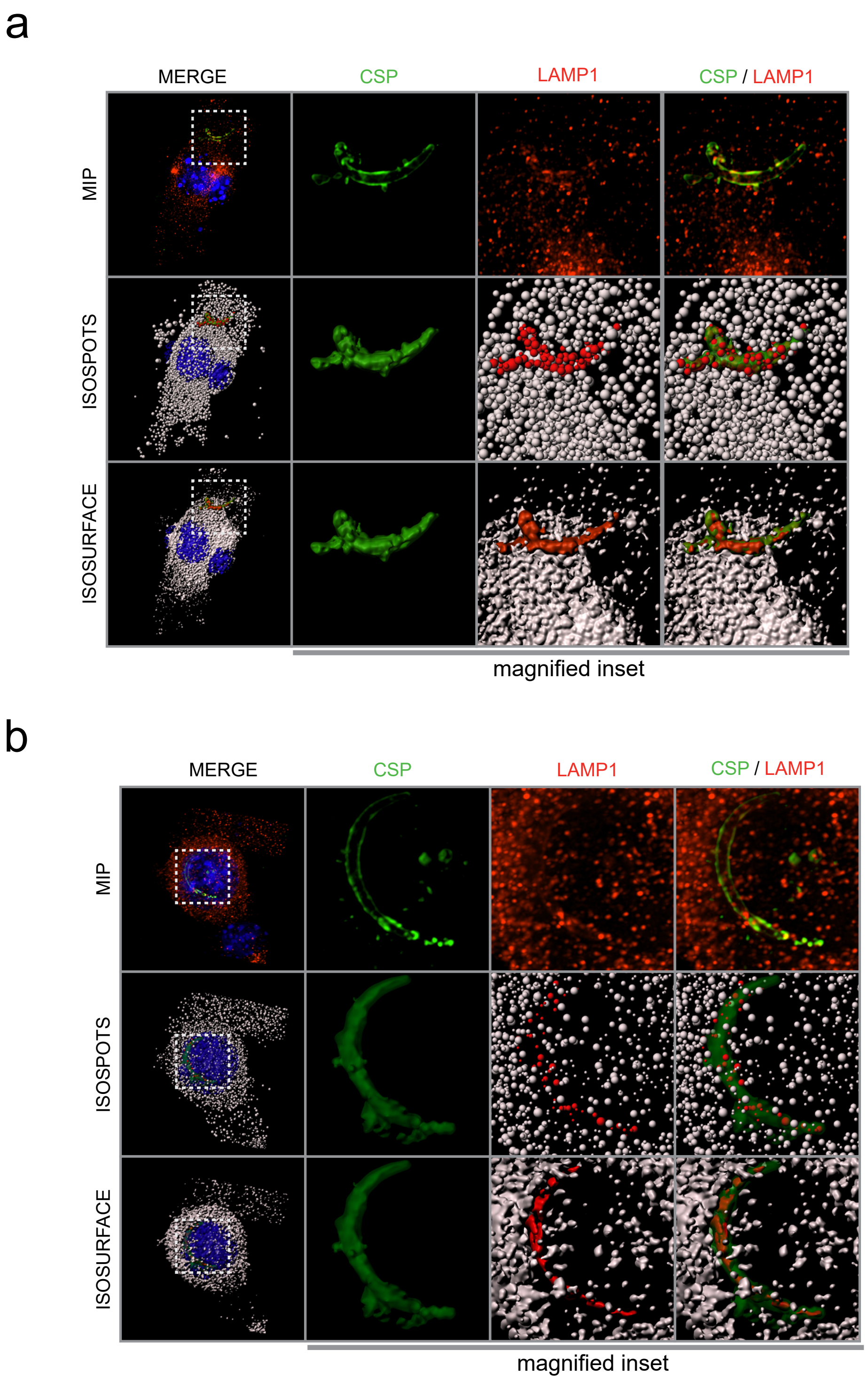

**Fig. S1. CT deficient or wild type *Plasmodium* sporozoites interacts with lysosomes in similar fashion.** Immunofluorescent microscopy of Hepa1-6 cells infected with SPECT2$^{-}$ *P. yoelii* sporozoites. Infection was assessed after **(a)** 5 or **(b)** 30 min and processed for fluorescence microscopy using DAPI (blue) for DNA, Phalloidin (white) for actin visualization, antibodies to LAMP1 (red) for LE/lysosomes and CSP (green) for parasites. Isospots for LE/lysosomes and isosurfaces for LE/lysosomes, parasites, host cell nucleus and plasma membrane were created using Imaris software and the LE/lysosomes interacting with parasites were identified by detecting overlap between isospots and the isosurface. Red spots represent LAMP1-positive structures co-localized with CSP. Magnified inset is 15 µm $x$ 15 µm.

**Supplementary Figure 2**

a

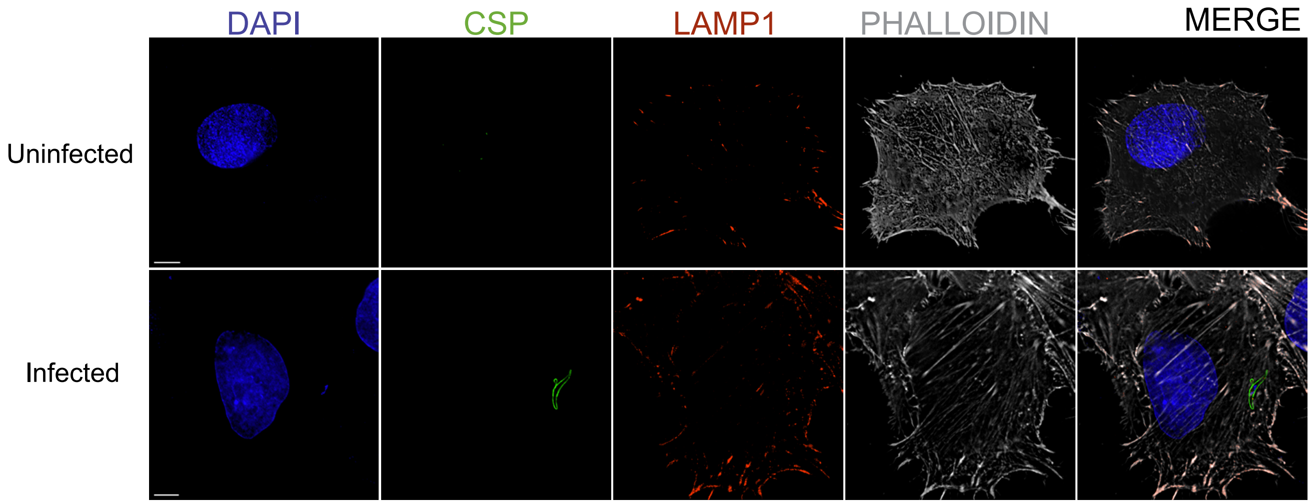


b

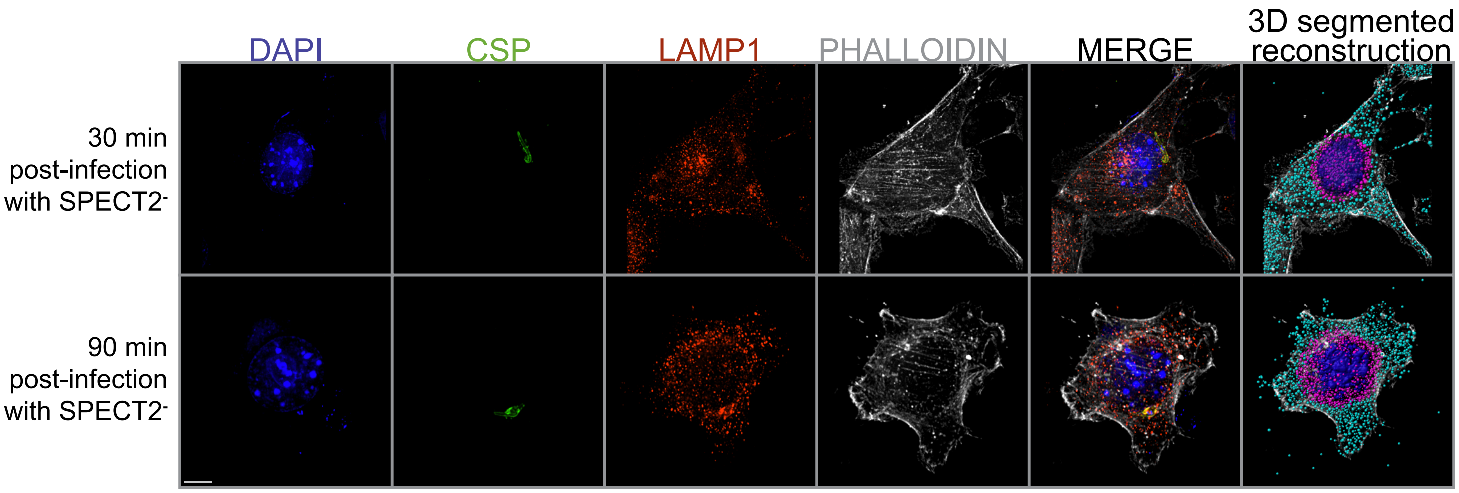

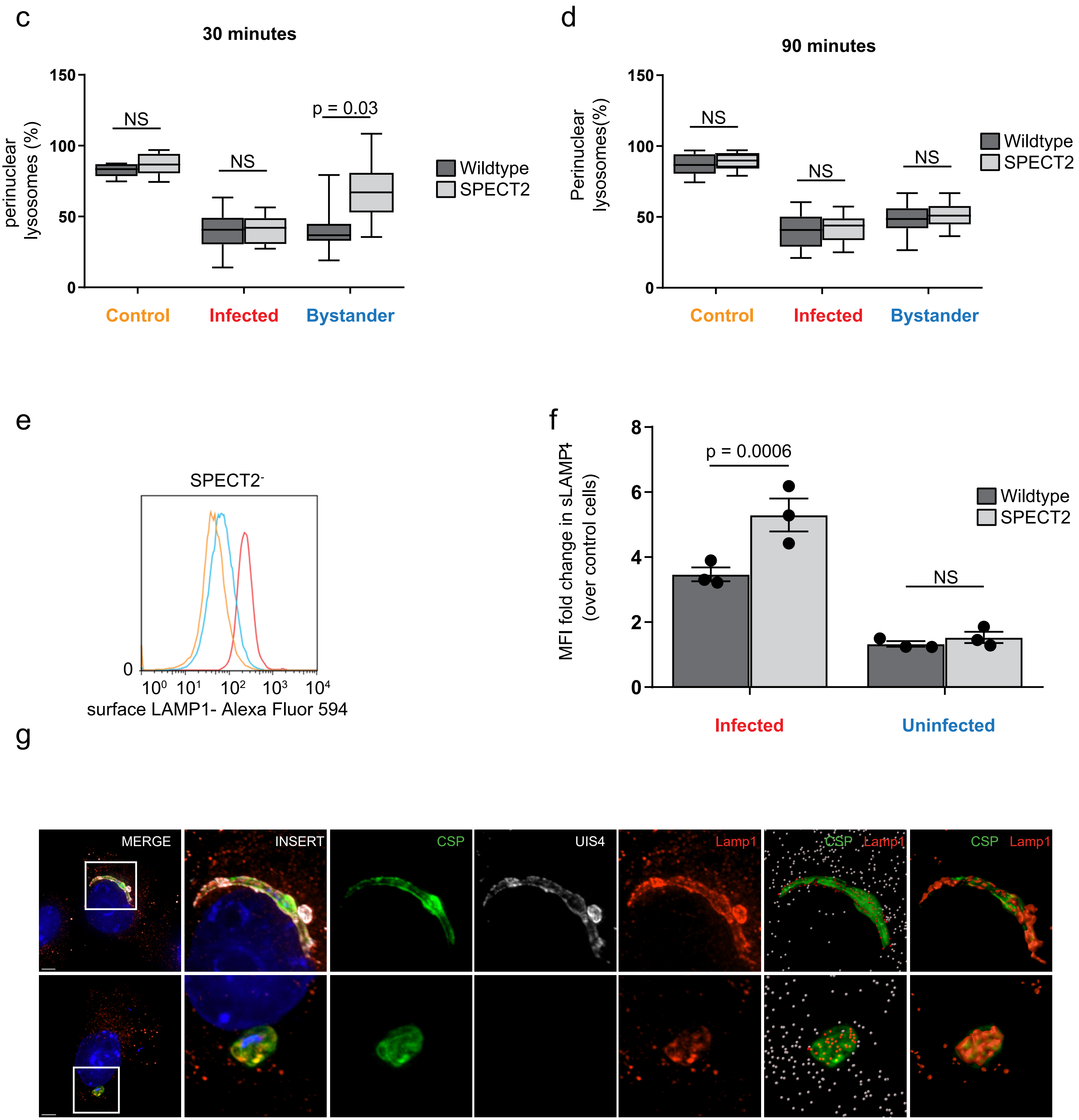
c
30 minutes
perinuclear lysosomes (%)
150
100
50
0
NS
NS
p = 0.03
Wildtype
SPECT2
Control
Infected
Bystander
d
90 minutes
Perinuclear lysosomes(%)
150
100
50
0
NS
NS
NS
Wildtype
SPECT2
Control
Infected
Bystander
e
SPECT2-
0
10⁰ 10¹ 10² 10³ 10⁴
surface LAMP1- Alexa Fluor 594
f
MFI fold change in sLAMP1+ (over control cells)
8
6
4
2
0
p = 0.0006
NS
Wildtype
SPECT2
Infected
Uninfected
g
MERGE
INSERT
CSP
UIS4
Lamp1
CSP Lamp1
CSP Lamp1

**Fig. S2. Sporozoite induced lysosome-plasma membrane fusion is independent of cell traversal. (a)** Hepa1-6 cells were infected with *P. yoelii* sporozoites and fixed after 90 min. Cells were stained with antibodies to LAMP1 prior to permeabilization and stained with DAPI (blue) for DNA, phalloidin (white) for actin visualization, antibodies to CSP (green) for parasites and displayed as maximum intensity projections. Bar = 5 µm. **(b)** Hepa1-6 cells were infected with SPECT2$^{-}$ sporozoites and fixed after 30 and 90 min. Cells were processed for 3D fluorescence microscopy using DAPI (blue) for DNA, phalloidin (white) for actin visualization, antibodies to LAMP1 (red) for LE/lysosomes and CSP (green) for parasites and displayed as maximum intensity projections. Scale bar = 5 µm. Images were obtained on a Deltavision fluorescent microscope and processed by Imaris software to construct isosurfaces (nuclei and parasite) and LAMP1-positive isospots (LE and lysosomes) by predefined algorithms for identification of surfaces and spots. Perinuclear isosurfaces were created by extrapolating the DAPI signal to define a perinuclear region where lysosomes could be differentially quantified. The isospots corresponding to total and perinuclear lysosomes were depicted in cyan and red, respectively. Bar = 5 µm. **(c and d)** Values represented in box and whiskers plot correspond to lysosomes from total and perinuclear area represented as mean ± SD of 25 different microscopic fields from three independent experiments. **(e)** Hepa1-6 cells were infected with SPECT2$^{-}$ *P. yoelii* sporozoites for 90 min and analyzed by flow cytometry using antibodies specific to LAMP1 and CSP. Surface LAMP1 was evaluated by staining cells prior to permeabilization, while total LAMP1 was evaluated by staining for LAMP1 after permeabilization. The histogram shows the distribution of surface LAMP1 from SPECT2$^{-}$ infected, uninfected and unexposed control cells from one of three independent experiments. **(f)** Surface LAMP1 levels were compared between uninfected and SPECT2$^{-}$ infected cells as a fold change over control cells. The bar graph depicts the mean ± the SD of three independent experiments. **(g)** Hepa1-6 cells were infected with *P. yoelii* sporozoites and fixed after 90 min. Cells were stained with DAPI (blue) for nuclei visualization, antibodies to LAMP1 for lysosomes, UIS4 (white) and CSP (green) for parasite detection and displayed as maximum intensity projections. Bar = 5 µm. Red spots represent LAMP1-positive structures co-localized with CSP. Magnified inset is 15 µm $x$ 15 µm.

#### Supplementary Figure 3

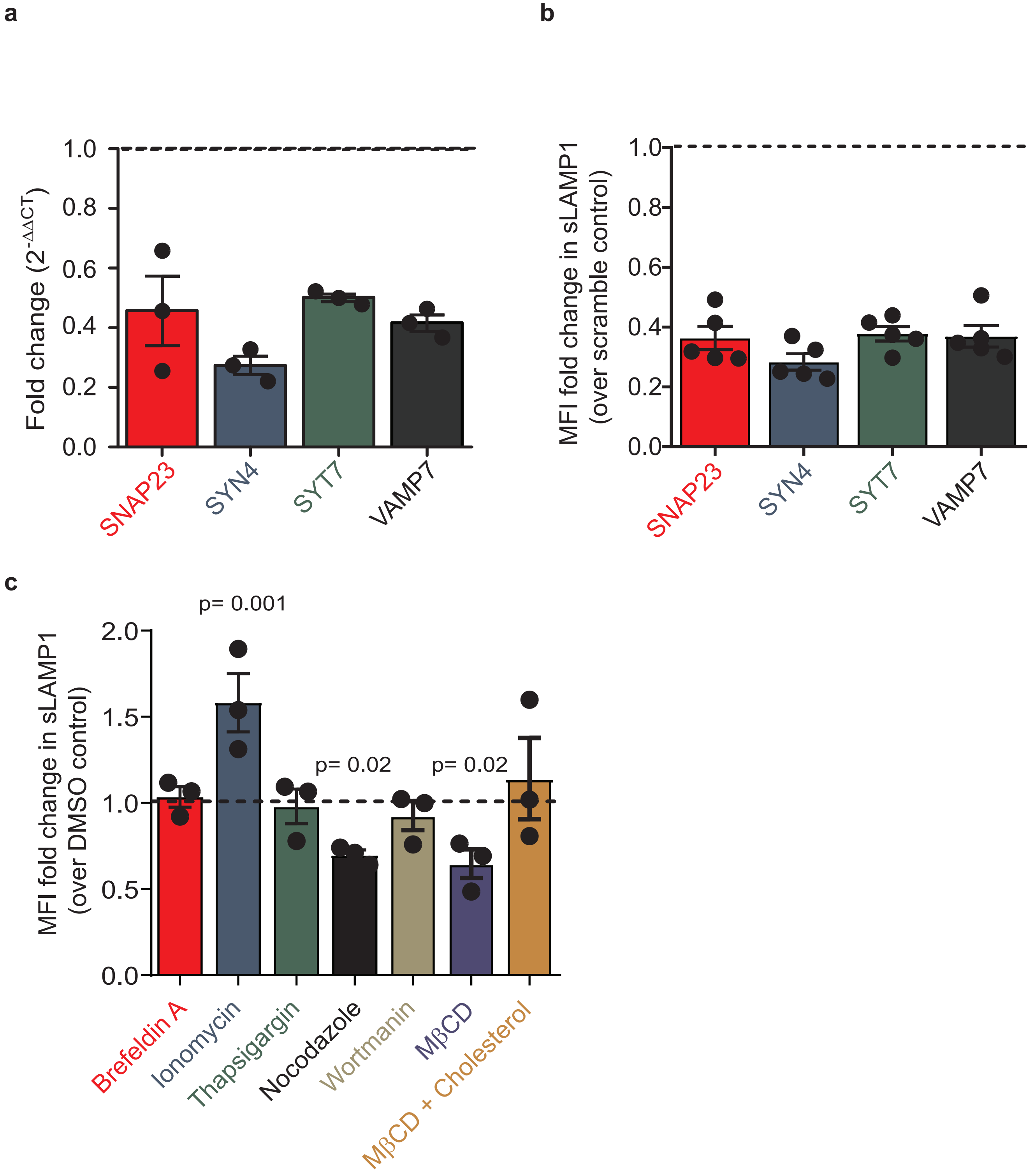

**Fig. S3. Selective knockdown of SNARE proteins can be achieved using lentivirus-mediated shRNA. (a)** Bar graph depicting relative gene knockdown compared to non-targeting control shRNA. Hepa 1-6 cells were transduced with lentivirus expressing shRNA constructs selectively targeting SNARE proteins, or a non-targeting control. Mean knockdown level was determined using qPCR. Values are normalized to non-targeting control which is indicated by solid black line. Data is mean ± the SD of 3 independent experiments. **(b)** Hepa1-6 cells were transduced with shRNA lentiviruses against SNAP23, SYN4, SYT7, or VAMP7 or a scrambled control and assessed for surface LAMP1 by flow cytometry. Surface LAMP1 levels were compared between scramble and specific knockdowns and expressed as a fold change in mean fluorescent intensity (MFI). The bar graph depicts the mean ± the SD of 5 independent experiments. **(c)** Hepa1-6 cells were incubated with or without 10 µM ionomycin, 400 nM thapsigargin, 10 µM brefeldin A, 10 µM nocodazole, 5 mM methyl-β-cyclodextrin (MβCD) for 15 min, fixed and assessed for surface LAMP1 by flow cytometry. For cholesterol replenishment, methyl-β-cyclodextrin (MβCD) treated cells were incubated with 1 mM cholesterol for 15 min prior to fixation. Surface LAMP1 levels were compared between DMSO vehicle control and specific treatments and expressed as a fold change in mean fluorescent intensity (MFI). The bar graph depicts the mean ± the SD of three independent experiments.

**Supplementary Figure 4**

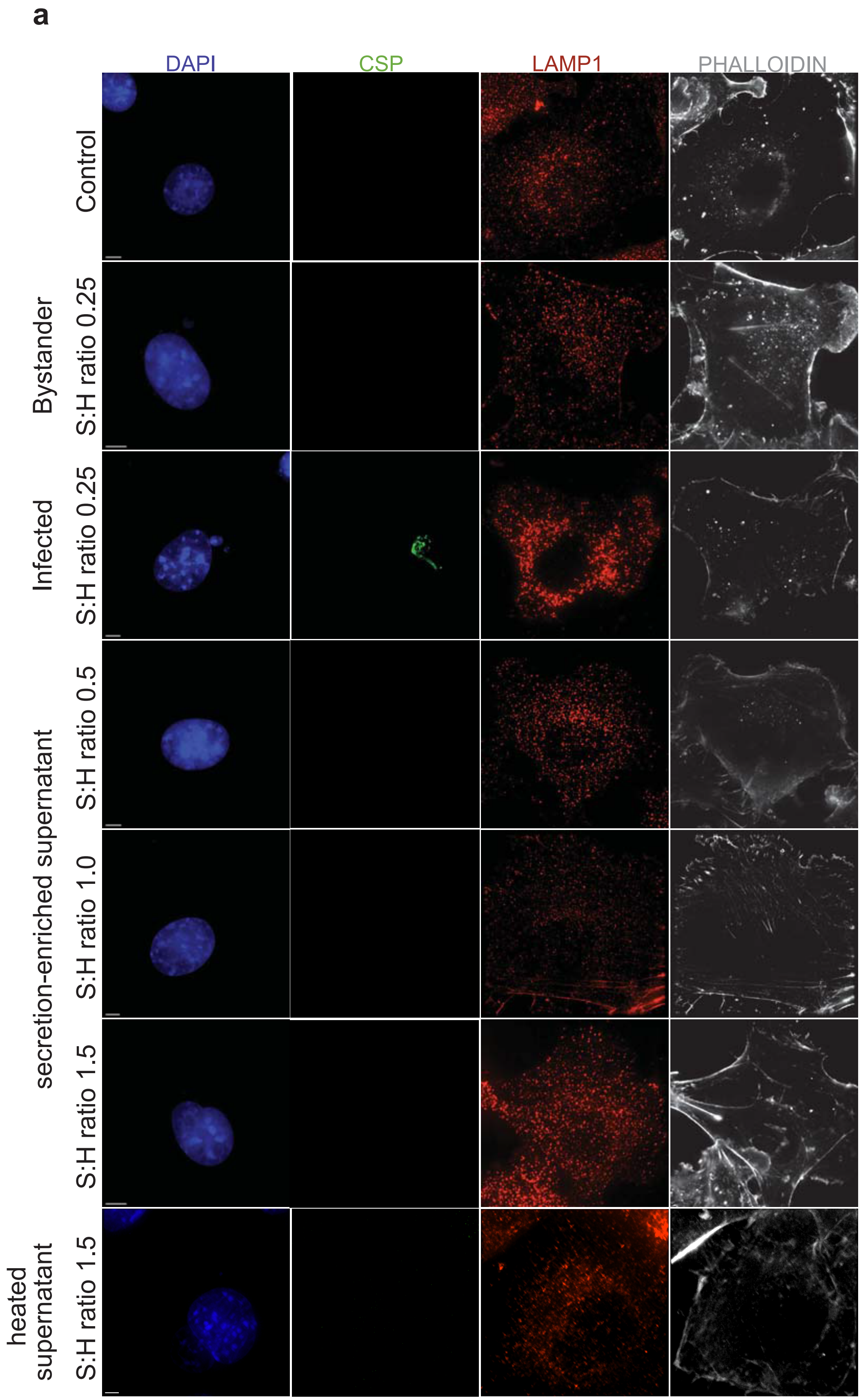

**Fig. S4. Sporozoite secreted factor contribute to relocalisation of lysosomes.** Hepa1-6 cells were infected with *P. yoelii* sporozoites or treated with FBS activated *P. yoelii* sporozoite secretion-enriched supernatants at different sporozoite:hepatocyte (S:H) ratios for 90 min. Cells were processed for 3D fluorescence microscopy using DAPI (blue) for DNA, phalloidin (white) for actin visualization, antibodies to LAMP1 (red) for LE/lysosomes and CSP (green) for parasites and displayed as maximum intensity projections. Bar = 5 µm.

**Supplementary Table 1**

Lysosomal trafficking modulators selected for the study.

| Inhibitor | Function | Reference |
|---|---|---|
| Ionomycin | ionophore, increases intracellular Ca++ and induces lysosome exocytosis. | (*3*) |
| Brefeldin A | redistributes LE/lysosomes towards periphery | (*4*, *5*) |
| Nocodazole | a microtubule-depolymerizing agent, prevents lysosome redistribution. | (*4*) |
| Thapsigargin | inhibitor of the sarco/endoplasmic reticulum Ca++ ATPase, elevates cytosolic Ca++ and promotes lysosome exocytosis. | (*6*) |
| Methyl-β-cyclodextrin | depletes membrane cholesterol and reduces LAMP1 levels on surface. | (*7*) |
| Wortmannin | PI3K inhibitor, inhibits various stages endocytic network. | (*8*) |
